# Nanopore Electrometry Resolves Peptide Charge Patterns beyond Ionic-Current Blockade

**DOI:** 10.64898/2026.07.20.739564

**Authors:** Pranjal Sur, Prabal K. Maiti, Manoj M. Varma

**Affiliations:** Centre for Nano Science and Engineering, Indian Institute of Science, Bengaluru-560012; Centre for Condensed Matter Theory, Department of Physics, Indian Institute of Science, Bengaluru-560012

**Keywords:** Solid-state nanopores, Protein sequencing, Single molecule sensing, Graphene, Molecular dynamics, Nanopore electrometry

## Abstract

Localized measurements of electric fields offer a promising route to expand the information content of nanopore-based single-molecule sensing beyond conventional ionic-current blockade. Here, using all-atom molecular dynamics simulations with virtual electric-field probes placed around a graphene nanopore, we show that the local electric-field captures the presence, and distribution of charged amino acids as the peptides translocate through the pore. These field signatures create reproducible peptide-specific fingerprints across independent translocation events and enable substantially improved discrimination between peptides compared with ionic-current traces obtained under the same simulation conditions. Our results suggest that localized nanopore electrometry can provide a complementary, information-rich readout of peptide charge order that is largely inaccessible to conventional current blockade-based measurement. This study establishes a simulation-guided framework for integrating nanoscale electrometry with nanopore platforms for future peptide and protein analysis.

---

Solid-state nanopores have emerged as a versatile single-molecule sensing platform^1^ over the past two decades, driven by advances in nanofabrication and characterization techniques. The success of biological nanopores in commercial DNA sequencing has established the power of nanopore-based molecular readout, most prominently through the Oxford Nanopore platform^2^ However, an analogous capability for native protein sequencing has not yet been achieved with either biological or solid-state nanopores. Recent studies have nevertheless demonstrated important progress toward this goal; biological nanopores have enabled sequence-resolved sensing of unfolded protein strands,^3^ while solid-state nanopores have recently been used for full-length protein classification.^4^ Solid-state nanopores are particularly attractive for such applications because they offer mechanical and chemical robustness, compatibility with diverse materials and harsh environments containing protein denaturing agents, and the possibility of scalable fabrication using semiconductor-processing workflows.^1^ These advantages have made them versatile single-molecule sensors across applications ranging from biomarker detection and single-molecule DNA analysis to plasmonic sensing, nanoreactors, and iontronic devices.^5^ For protein and peptide analysis, solid-state nanopores are also useful probes of molecular structure, conformation, and translocation dynamics.^6–9^ At the same time, their use for sequence-level analysis remains limited by the rapid field-driven motion of analytes and by the spatial averaging inherent to conventional nanopore readout^5,10,11^. To improve spatial resolution, ultrathin membranes based on materials such as silicon nitride, MoS_2_, and graphene have been explored, with graphene representing an extreme limit in membrane thickness.^5,10^

While ultrathin membranes improve spatial confinement and can enhance the geometric resolution of nanopore sensing, the information obtained from a translocation event is still ultimately determined by the physical quantity being measured. In practice, nanopore experiments have relied predominantly on ionic-current blockade as the primary readout. This signal has enabled remarkable progress, but it may not be optimally sensitive to molecular features needed for sequence-level discrimination, such as the local distribution of charges and chemically distinct residues along a translocating peptide. In ionic current based measurement, an applied voltage drives ions through an electrolyte-filled nanopore and the passage of an analyte transiently obstructs ion transport producing a current drop. The blockade amplitude represents the strength of this obstruction, while the event duration reports the dwell time of the molecule in the pore. This simple and powerful measurement has enabled a broad range of nanopore applications, but it also has fundamental limitations for peptide and protein analysis. First, fast translocation in solid-state nanopores can push molecular events beyond the effective bandwidth of standard current amplifiers, creating a trade-off between temporal resolution and signal-to-noise ratio. Second, ionic current is a transport observable that is primarily governed by the analyte’s excluded volume and its perturbation of ion flow.^12,13^ It can therefore encode changes in molecular size, shape,^13,14^ and net charge,^9,15^ but it is not intrinsically capable to resolve the positional distribution of charges along a translocating peptide. This limitation is especially important for proteins, where sequence information is encoded not only in overall size or charge but also in the order and local placement of chemically distinct residues. This motivates the exploration of complementary readout modalities that can potentially access molecular information not readily captured by ionic current alone.

In our previous work, we introduced nanopore electrometry as a complementary readout strategy in which molecular translocation is detected through changes in the local electric field rather than through ionic-current blockade.^16^ That study established the physical basis for using localized field measurements in nanopores and highlighted potential advantages such as sensitivity to molecular charge, spatially distributed sensing, and self-referenced detection. However, the question of whether such a readout can capture chemically specific information from a translocating biomolecule remains open. In the present study, we address this question by examining whether localized electric-field traces can encode peptide charge-order fingerprints during nanopore translocation. This is particularly important for peptide and protein analysis, where the relevant information is not contained only in molecular size or net charge, but also in the local placement and ordering of charged residues along the chain. A possible route to experimental realization of nanopore electrometry is provided by nitrogen-vacancy (NV) centers in diamond, which are optically addressable solid-state spin defects capable of nanoscale quantum sensing at room temperature.^17^ Although NV centers are more widely used for magnetometry, electric-field sensing with single diamond spins has been demonstrated,^18^ including measurements of internal fields in semiconductor devices.^19^ NV-based electric-field imaging has also been achieved using shallow centers,^20^ and recent work has extended electric-field sensing to liquid electrolytes using nanodiamonds.^21,22^ These developments suggest a plausible experimental pathway for localized electric-field readout near nanopores.

To motivate the use of local electric-field signals for peptide analysis, we first asked whether charged residues alone carry substantial discriminative information in protein sequences. We analyzed the human proteome from UniProt^23^, restricting the dataset to manually curated Swiss-Prot entries and selecting one representative protein sequence per gene. For each sequence, we considered only the four charged amino acids expected to contribute most strongly to local electric-field modulation; namely, Aspartate/D and Glutamate/E, which are negatively charged, and Lysine/K and Arginine/R, which are positively charged. Using only the counts of these four residues, 93.5% of the analyzed protein entries could be distinguished from all others in the dataset (SI section 2). When the sequential order of these charged residues was also considered, the distinguishability increased to 98.2% (Figure 1a). This analysis highlights that charged-residue composition and ordering contain substantial molecular information that could be exploited for charge-order-based peptide fingerprinting using localized electric-field readout. We next selected a small set of naturally occurring human peptides to test whether such charged-residue patterns can be reflected in local electric-field traces during nanopore translocation. For this purpose, we filtered the UniProt entries to retain experimentally supported sequences, corresponding to protein existence level 1, with lengths between 15 and 20 residues. From this set, we selected four 16-residue peptides containing different numbers, signs, and arrangements of D, E, K, and R residues (Figure 1b). These peptides provide a compact model system for examining whether localized electric-field measurements at a graphene nanopore can distinguish peptide-specific charge-order fingerprints.

**Figure 1:**
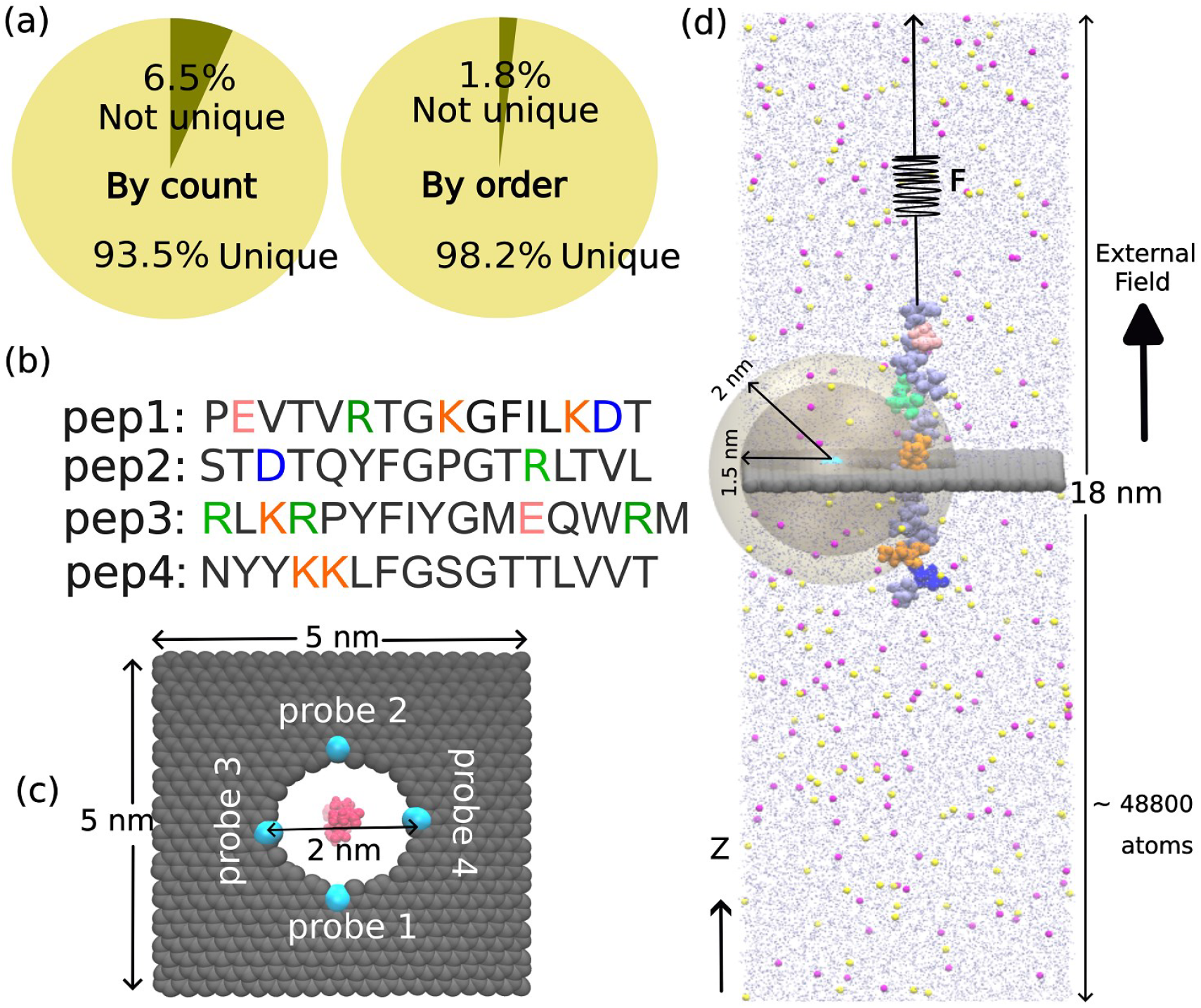
(a) Charged-residue-based distinguishability of human protein sequences using the four charged amino acids D, E, K, and R. The left chart shows distinguishability based only on the counts of these residues, while the right chart includes their sequential order. (b) Four naturally occurring peptide sequences selected for molecular dynamics simulations, with charged residues highlighted: D/E in blue/pink tones for negatively charged residues and K/R in orange/green tones for positively charged residues. (c) Top view of the monolayer graphene membrane containing a 2 nm diameter nanopore. Cyan spheres indicate the four virtual probe positions at which time-dependent electric fields were calculated. (d) Simulation setup for steered peptide translocation through the graphene nanopore in electrolyte under an applied electric field. The peptide is pulled along the z-direction through the pore while local electric-field signals are recorded at the probe positions. Two sensing regions defined by radial cutoffs 1.5 nm and 2 nm around each probe, were used. The molecules residing within the sensing region contributed to the cut-off specific instantaneous electric field.

We then used steered molecular dynamics simulations (SI section 1) to translocate the selected peptides through a monolayer graphene nanopore under an applied bias of 500 mV (Figure 1d). The graphene membrane and peptide were immersed in 0.5 M KCl, and four positions along the nanopore edge were selected as virtual electric-field probes (Figure 1c). At each probe, the instantaneous electric field was calculated throughout the translocation trajectory. The electric field was evaluated using molecules within a finite sensing region (Figure 1d) surrounding the probe. This work focuses on the field variations associated with nearby peptide, ion, and water dynamics, which are expected to dominate the translocation-induced signal.

The raw traces exhibit substantial thermal and electrostatic fluctuations (Figure S1), and the noise appears to be qualitatively similar for two different sensing regions with 1.5 nm and 2 nm cutoff respectively. We therefore applied the denoising procedure (SI section 1) to compare peptide-dependent signal features.

Figure 2 (a, b) shows representative traces of the z-component of the electric field for pep1. Representative traces for other peptides are shown in Figure S2, S3. The field modulations occur at times that coincide with the passage of charged residues or charged-residue clusters near the probe region, suggesting that local electric-field readout is sensitive to the charge distribution along the translocating peptide. In contrast, the corresponding ionic-current trace (Figure 2c) show weaker correspondence with the passage of charged residues through the pore under the same simulation conditions. The qualitative behavior of a field signature does not change significantly when the sensing region is increased from 1.5 nm to 2 nm (Figure S2, S3), indicating that the dominant contributions originate from the local environment around the probe.

**Figure 2:**
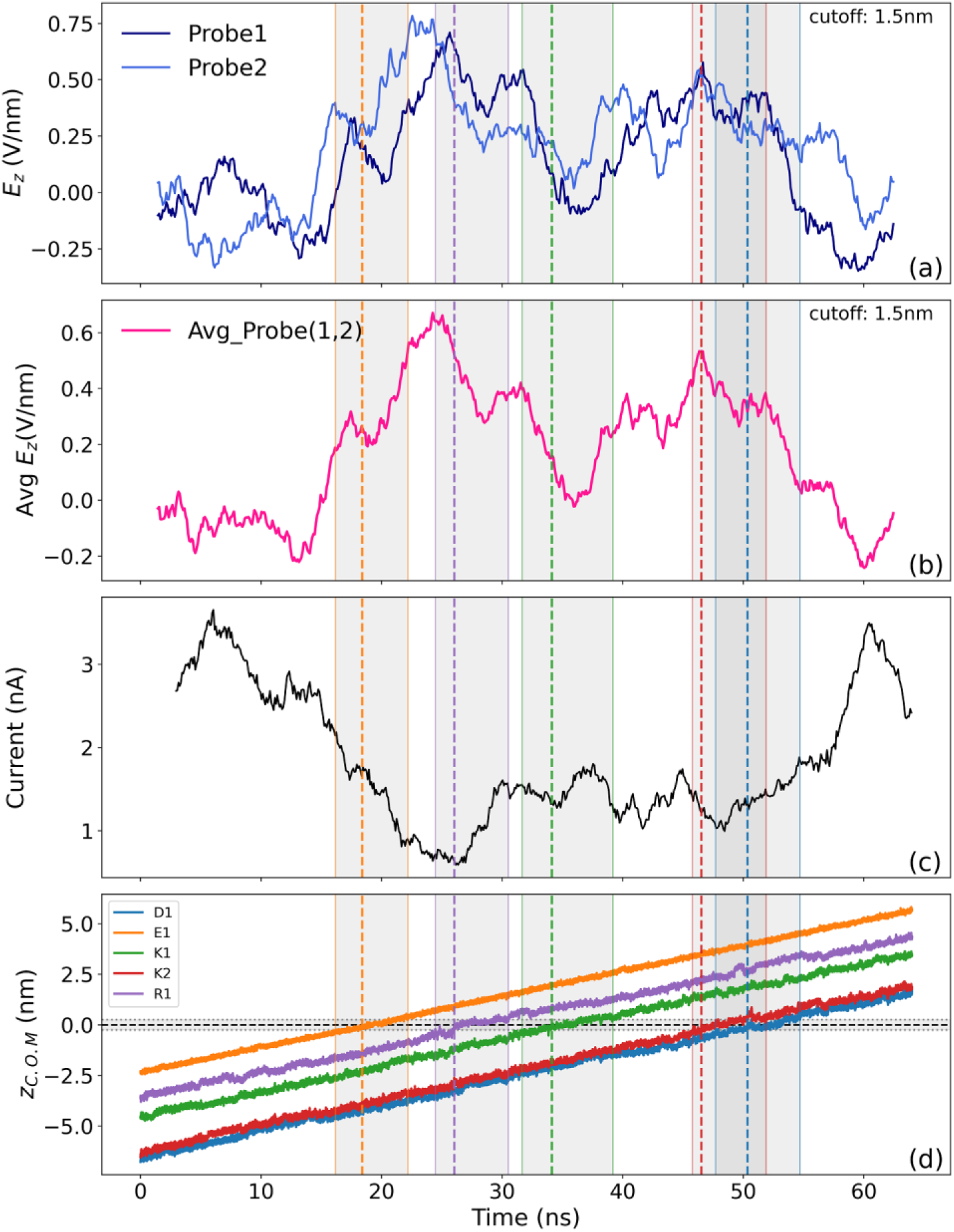
Time dependent signals while pep1 translocates through the pore. (a) Electric field traces captured by different probes. (b) Electric field trace after averaging over data obtained from two probes. (c) Ionic current signature. (d) Position of the charged residues of pep1 with respect to the pore. The gray bands indicate their passage through the pore within 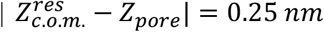. The band edges are color matched with the residues. The dotted vertical lines represent 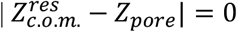. The residue label, for example, K2 represents residue K that is arriving at the pore for the second time, as per the sequence of the peptide.

The local nature of the electric-field measurement is further reflected in the probe-to-probe variation of the signals (Figure 2a, Figure 3). Probes placed at different positions along the nanopore boundary will not report identical field traces, because the measured field depends on the instantaneous configuration of the charged side chains and the surrounding ionic environment sampled at each probe. In the present simulations the backbone C_α_ atoms retain lateral position restraints in the X and Y directions during translocation (Section 1, SI), so that this variability originates from the motion of the unrestrained side chains, ions, and water rather than from lateral displacement of the peptide backbone. Such probe-specific variation is not necessarily a limitation. On the contrary, it suggests that spatially distributed electric-field probes could increase the information content of nanopore translocation measurements by simultaneously sampling different electrostatic environments around the translocating molecule. (Figure 3, Figure S7, S8).

**Figure 3:**
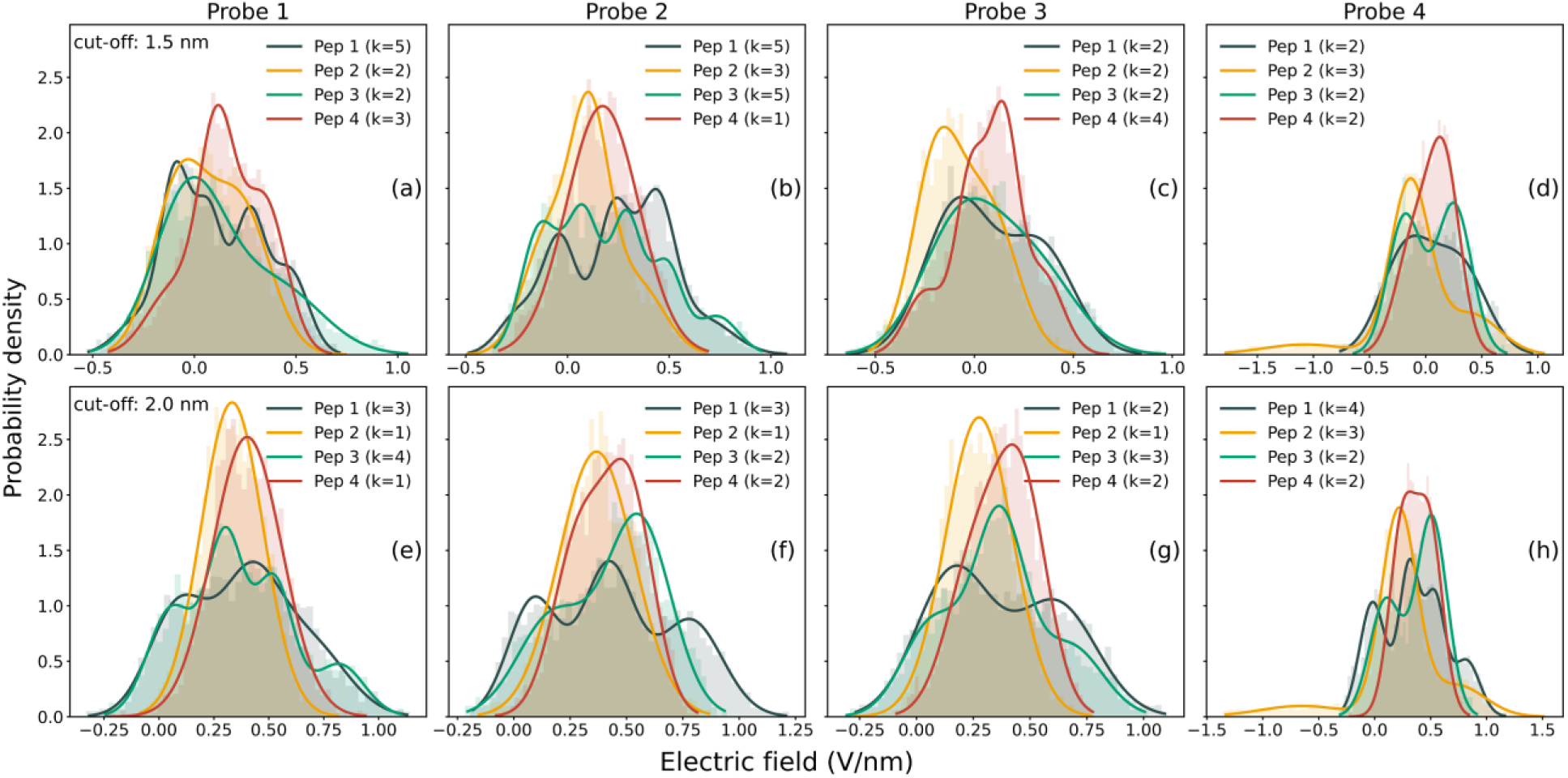
Gaussian Mixture Model (GMM) fit of histograms constructed from the z component of the electric field. The number of the gaussians describing the best fit is denoted as k in the legend of each subplot. (a-d) correspond to the sensing region within cutoff 1.5 nm whereas (e-h) corresponds to the sensing region within 2 nm cutoff. Each of the four column shows data from four different probes.

To characterize the distribution of z component of the electric field associated with each peptide, probe and sensing configuration, data from ten independent translocation trajectories were combined into a single ensemble to increase the statistical sampling of the signal distribution. The resulting histograms constructed from the ensembles were fitted with Gaussian Mixture Models (GMM). Models containing one to five gaussian components were evaluated using the Bayesian Information Criterion (BIC)^24^ and the model yielding the lowest BIC score was selected as the best representation of the underlying signal distribution.

The distributions, shown in Figure 3 reveal pronounced differences in modal complexity that depend on peptide charge content, probe position and sensing volume. At the smaller sensing volume with cut-off radius 1.5nm (Figure 3 a-d), several peptide-probe combinations exhibit highly structured distributions requiring up to five gaussian components for optimal representation. In particular, pep1 and pep3, each containing five charged residues, generally display more complex distributions than pep2 and pep4 which contain two charged residues. This trend suggests that increasing charge content increases the diversity of electrostatic environments sampled during translocation. However, the relationship is not purely determined by the number of charged residues. The differences in modal structure indicate that the probe response depends not only on charge count but also on the spatial arrangement of charge along the peptide. For example, in Figure 3 b, f, the distribution for pep1 and pep3 is visibly different, where they show different number of distinct modes, respectively, although they contain the same number of charged residues. Increasing the sensing cut-off from 1.5 to 2 nm (Figure 3 e-h) reduces the complexity of the electric field distributions, leading to fewer (SI section 5) distinguishable gaussian components. Hence, the 1.5 nm sensing radius provides greater sensitivity to perturbations in the local electrostatic environment compared to the 2 nm sensing radius. The four sensing probes exhibit distinct signal distributions for the same peptide, for both the sensing volume. This suggests that probes positioned at different locations around the nanopore capture partially independent information on surrounding electrostatic atmosphere that is spatially heterogeneous and provide complementarity rather than redundant measurements.

Ionic current distributions and corresponding GMM fits shown in Figure 4 were constructed following the same workflow used for electric field analysis in Figure 3. Although the ionic current distributions also exhibit multimodal behavior, their modal structures differ from that observed for the electric field traces, especially for pep1 and pep3. For example, if the green curve of pep3 in Figure 3b (field) and in Figure 4 (current) are compared, it is noticeable that the neighboring electric field distribution peaks appear better resolved than ionic current distribution peaks. To quantify the degree of separation between adjacent modes in the fitted GMM models (Figure 3 and 4), and to compare among the field and current distributions, a peak-resolution analysis (SI section 5) was performed using the fitted model parameters. The results (Figure S4, S5, and S6) suggest that the field distributions are better resolved than the ionic current.

**Figure 4:**
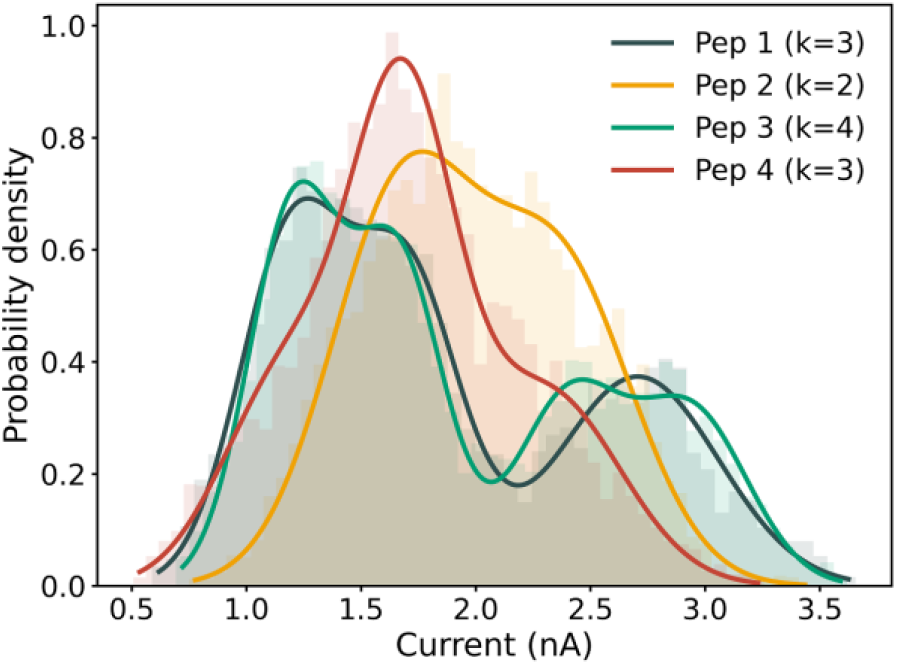
Gaussian Mixture Model (GMM) fit of histograms constructed from the ionic current traces. The number of the gaussians describing the best fit is denoted as k in the legend.

We next asked whether these peptide-dependent field features are statistically reproducible across independent translocation events. As mentioned before, we performed ten independent replica simulations for each peptide and recorded the time-dependent electric-field and ionic-current signals for every trajectory. Signal similarity was quantified using dynamic time warping (DTW), which measures shape-based similarity between time series while allowing for differences in event duration and local time alignment^25^. Each trajectory was labelled by peptide and replica number; for example, P1R1 denotes replica 1 of pep1. Because the peptide is advanced through the pore, starting from a common equilibrated configuration, by pulling a terminal residue at a fixed velocity while the C_α_ atoms were positions restrained in X and Y direction, all replicas of a given peptide traverse the pore at the same nominal rate and share a common translocation schedule. The variation among them arises from the thermal motion of the unrestrained side chains, the limited flexibility of the backbone permitted by the restraints, and the surrounding ionic and solvent environment. This has two consequences for the analysis that follows. First, translocation speed and dwell time are eliminated as confounding variables, so that the discrimination between peptides reported here reflects the charge-order-dependent structure of the field rather than differences in translocation kinetics. This is a meaningful control, as dwell time is a dominant discriminator in ionic-current sensing. Second, the imposed kinematics and the shared initial configuration suppress the dispersion in translocation time and pathway that would arise under a purely voltage-driven process, so that the within-peptide variability of the field traces is expected to be smaller than under unbiased translocation. The class separability reported below should accordingly be regarded as an upper bound attainable under controlled kinematics rather than as a direct estimate of experimental performance. The dynamic time warping distance used to compare traces is furthermore invariant to local time rescaling, so that the resulting clustering reflects the shape and ordering of the field modulations rather than their absolute timing. Figure 5 compares the resulting DTW distance matrices from the z-component of the electric field evaluated using two different sensing radii and from the ionic current. The DTW distances were computed between pairs of traces from the corresponding pair of replica trajectories. DTW distance matrices for the electric field traces were generated not only for individual probes (Figure S7) but also for signals obtained by averaging all probes (Figure S8) and different probe combinations. A given electric field trace could be constructed from any combinations of the four probes. After evaluating all possible probe combinations using hierarchical clustering, the average signal obtained from probes 1,3, and 4 yielded the highest cluster separation. Consequently, the electric-field analysis shown in Figure 5, is subjected to an average of the signals from probes 1, 3, and 4.

**Figure 5:**
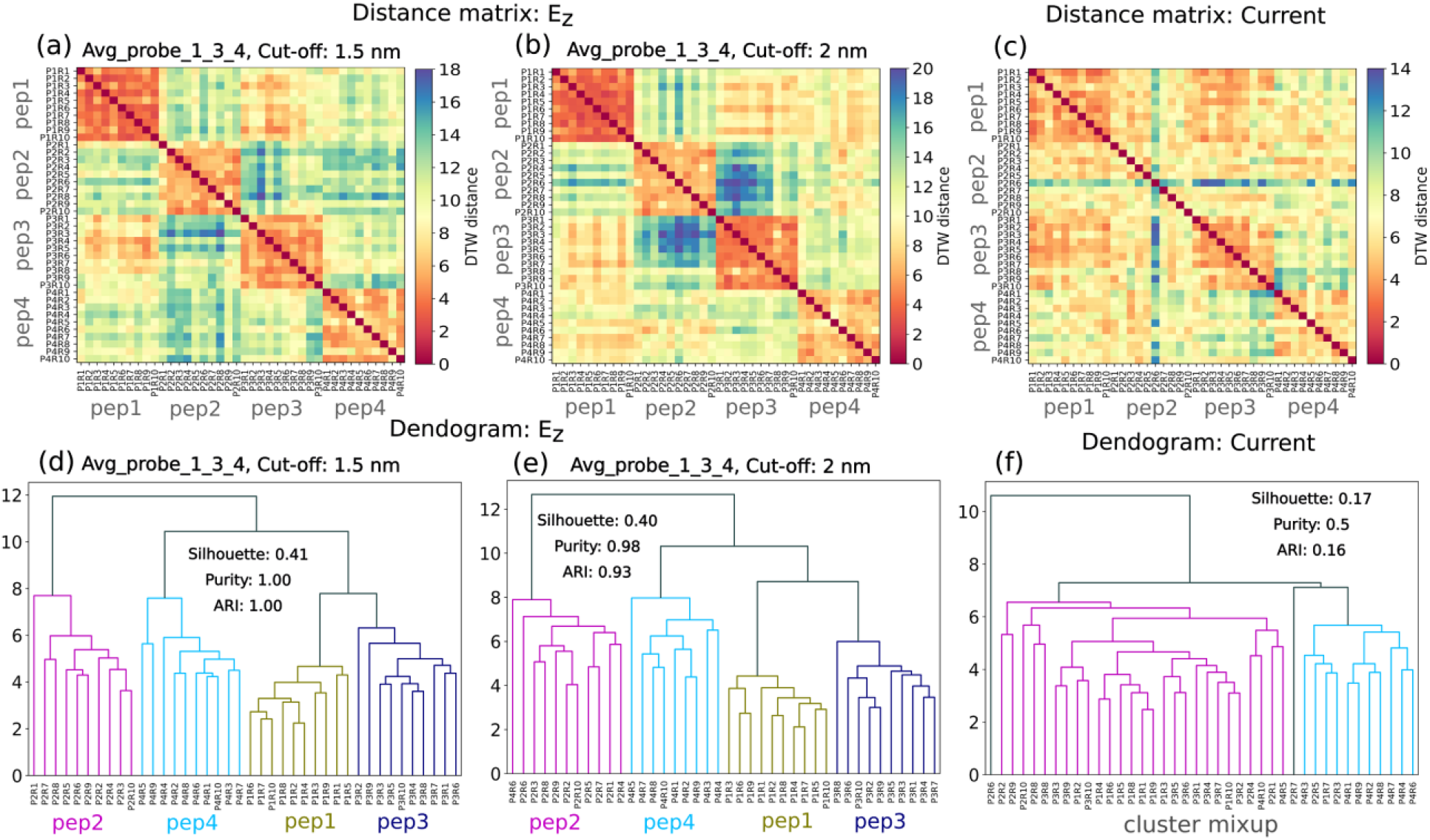
Dynamic time warping (DTW) distance matrices computed from ten independent translocation replicas of each peptide using the averaged z-component of the electric field from probes 1, 3, and 4 within sensing volume corresponding to radial cut-off of (a) 1.5nm and (b) 2nm. (c) DTW distance matrix of the corresponding ionic-current traces. Hierarchical clustering of the same DTW distances for (d) electric field within 1.5nm sensing radius, (e) electric field within 2nm sensing radius and (f) ionic current.

The electric-field distance matrix shows a clear block-diagonal structure (Figure 5 a, b), indicating that replicas of the same peptide are more similar to one another than to trajectories from other peptides. This feature is present for both the sensing radii of 1.5nm and 2nm. This peptide-wise organization is much weaker in the ionic current-based distance matrix, where inter-peptide mixing is more apparent (Figure 5c). Hierarchical clustering under a four-cluster constraint further emphasizes this difference. Electric-field signals produce well-separated clusters corresponding to the four peptide identities (Figure 5d, e), whereas current-based clustering shows substantial overlap between peptide classes (Figure 5f). Quantitatively, the electric-field feature space shows strong agreement with the known peptide labels, with the 95% confidence intervals for Rand indices, purity, and silhouette coefficient being [0.63, 1.00], [0.75, 1.00], and [0.32, 0.43], respectively, for the 1.5 nm case, and [0.85, 1.00], [0.94, 1.00], and [0.37, 0.41], respectively for the 2 nm case. These confidence intervals were obtained by subsampling over replica trajectories (SI section 6). In comparison, the corresponding ranges were [0.12, 0.27], [0.47, 0.56], and [0.13, 0.20], respectively for ionic-current based clustering showing substantially poorer class separation. The confidence intervals for the field- and current-based clustering do not overlap at either sensing volume indicating that the superior class separability of the electric-field readout is statistically resolved at the level of sampling afforded by the available replicas. It should be noted, however, that the perfect index obtained at the 1.5 nm sensing volume is the more sensitive to the particular set of replicas: under removal of individual replicates its adjusted Rand index ranges over [0.63, 1.00], whereas the corresponding value at 2 nm remains within [0.85, 1.00], the median being unchanged in both cases. This greater stability of the larger sensing volume is consistent with the reduced sensitivity but higher reproducibility of the 2 nm signal, and the perfect separation reported at 1.5 nm should accordingly be regarded as a best case attainable under the controlled kinematics rather than a guaranteed outcome.

Taken together, these results show that localized electric-field readout provides a molecularly informative signal during peptide translocation through a graphene nanopore. Unlike ionic current, which primarily reflects ion transport through the pore and shows limited sensitivity to the positional arrangement of charged residues, the local electric field retains features associated with the charge distribution of the translocating peptide. This difference is evident in the denoised traces, where electric-field signals produce well-separated peptide-specific clusters. These findings support the central premise of nanopore electrometry: measuring local electric fields can provide a complementary information channel that is not readily accessible from ionic-current blockade alone. Experimental realization of this approach will require advances in nanoscale electric-field sensing, including control over sensor placement, sensitivity, bandwidth, and operation in electrolyte environments. A specific practical constraint concerns the spatial and temporal resolution attainable by a physical field sensor. Spatially, an optically addressable spin sensor such as a nitrogen-vacancy center reports the field at a finite standoff and averages it over a finite sensing volume, so lateral features of the charge distribution finer than this scale are attenuated. The present simulations provide a first, if idealized, probe of this dependence: the comparison between the 1.5 nm and 2 nm sensing volumes shows that the charge-order signatures degrade gradually rather than abruptly as the sensing volume is enlarged (Figures 3 and 5), indicating a degree of tolerance to finite spatial resolution while also showing that the smaller sensing volume preserves more of the discriminating structure. Temporally, field sensing is subject to a bandwidth–sensitivity trade-off analogous to that of ionic-current amplifiers, in which the averaging required to reach adequate field sensitivity sets a lower bound on the resolvable event duration. We therefore emphasize that the results presented here define the resolution regime, both spatial and temporal, that a physical sensor would need to reach, rather than demonstrating that present sensors already operate within it. Whether the charge-order signatures survive a realistic readout combining finite standoff, a fixed sensing axis, and coherence-time-limited averaging remains an open question that the present work is intended to motivate rather than settle. Nevertheless, ongoing progress in localized electrometry, including NV-center-based sensing, suggests a plausible route toward integrating electric-field readout with nanopore platforms. Such hybrid approaches could enable sequence-informed peptide fingerprinting and, more broadly, enrich the information available from single-molecule nanopore measurements.

## Supporting information

Supplementary information for the main manuscript

## Data availability

The data that support the findings of this study are available from the corresponding authors upon reasonable request.

## Acknowledgements

This work was supported by the Scientific and Useful Profound Research Advancement (SUPRA) Program of the Science Engineering Research Board (SERB) under Grant SPR/2021/000275 and the Scheme for Translational and Advanced Research in Science (STARS) under Grant MoE-STARS/STARS-2/2023-0077. P. S. acknowledges the PhD fellowship support from DST, India.

