## Supplementary information for the main manuscript for "Nanopore Electrometry Resolves Peptide Charge Patterns beyond Ionic-Current Blockade"

#### 1. Methods

All-atom molecular dynamics simulations were performed using GROMACS[1]. Inter-and-intra-molecular interactions were described using the CHARMM36m[2] force field. Graphene was modelled using the CA atom type from the same force field, treated as a charge-neutral membrane. Our water model was the CHARMM modified TIP3P[3] and we use ion-interaction parameters developed by Beglov and Roux[4]. To maintain the rigidity of the water molecules we used Settle[5], while LINCS[6] was implemented for hydrogen bond constraints. Our system contained a 500mM KCl solution. An external field of 0.0277 V/nm was applied along +ve Z creating a 500mV bias across 18nm long simulation box. Following a steepest-descent energy minimization, a two-equilibration protocol was applied. First an NVT equilibration was carried out for 5ns using a velocity-rescaling thermostat[7] at 300K with a coupling constant of 0.1ps. This was followed by a 5ns NPT equilibration using an isotropic Berendsen barostat[8] (coupling constant=2ps) and same temperature control as describe earlier. This ensures reaching equilibrium density at 1 bar reference pressure. 64 ns long production simulations were subsequently performed under NVT conditions with the velocity-rescaling thermostat, using a time constant of 1ps. Constant velocity pulling of the peptide was performed with a pulling rate of 1.25 Å/ps and a spring constant of 1500 kJ/mol/nm$^2$. Steered molecular dynamics was employed to obtain complete translocation events within the accessible simulation time. Under the applied bias alone, spontaneous capture and threading of a peptide into the nanopore could take prohibitively long time. So constant velocity pulling of the peptide is required to obtain a full translocation within a fixed simulation window to manage computational cost. For each peptide, ten replica trajectories were generated from a common equilibrated configuration. The dispersion among replicas therefore qualitatively represents a lower bound on the configurational variability that would be encountered under unbiased translocation. Long-range electrostatic interactions were computed using the Particle Mesh Ewald (PME) method[9]. A cutoff of 12Å was used for short-range Lennard-Jones and Coulomb interactions. A switching function was applied between an inner cutoff (10 Å) and outer cutoff (12 Å) for the LJ potential, ensuring smooth decay of both energy and forces to zero, thereby avoiding artifacts from force discontinuities. The system temperature was maintained at 300K throughout. We used an integration time step of 2 fs and trajectories were recorded every 2ps.

Position restraints with force constant of 1000 kJ/mol/nm$^2$ were applied to the $C_\alpha$ atoms of each residue. During minimization, restraints were removed, during equilibration they were applied in all three directions. In production run, we removed the restraints in Z direction. Strong position restraints (50000 kJ/ mol/nm$^2$) were present on graphene atoms in all stages of the simulations. Periodic boundary conditions were applied in all three directions. To denoise the raw electric field and current, we block averaged the signals over 100ps window followed by a moving average of 3 ns window. VMD[10–12] was used for all visualization purposes. Linear peptide chains were constructed using the tleap module in AmberTools[13]. Electric fields were calculated using the TUPÃ[14] python module, which leverages the selection capabilities of MDAnalysis[15,16] to compute electric fields as real space superpositions of coulomb contributions at user defined probe points, including contributions from periodic images within a specified cutoff distance. The work[14] introducing TUPÃ demonstrated that electrostatic contributions beyond a 15 Å cutoff are negligible for such localized field calculations. Accordingly, we chose a 15 Å cutoff around the probe points. Contributions from peptide, water molecules and ions residing within the cutoff boundary were included in the electric field evaluation. Additionally, we also calculate the electric field with a 20 Å cutoff for a qualitative comparison of the key results. The time dependent current signal was calculated using,

$$I\left(t+\frac{1}{2}\Delta t\right) = \frac{\sum_{i=1}^{N} q_i\,[z_i(t+\Delta t) - z_i(t)]}{\Delta t \times L_z}$$

Where, $q_i$ and $z_i$ are respectively the charge and position of $i^{th}$ ion, N is the total number of ions at time t, $\Delta t$ is the interval between each frame, and $L_z$ denotes the simulation box length in Z. A cylindrical region of 2nm height centered around the pore was considered for current calculations.

#### 2. Estimating protein sequence uniqueness

For each protein sequence, a reduced sequence was generated by retaining only the charged amino acids ASP (D), GLU (E), LYS (K), and ARG (R). This reduced sequence preserves the order of occurrence of charged residues along the primary sequence. Protein uniqueness was evaluated using two metrics:

1. Count-based uniqueness: For each protein, a four-element composition vector was calculated containing the total numbers of D, E, K, and R residues. Proteins sharing the same composition vector ($N_D$, $N_E$, $N_K$, $N_R$) were grouped together. A protein was considered unique by count if no other protein in the proteome possessed the same charged-residue composition vector.

2. Sequence-order uniqueness: The reduced charged-residue sequence itself was used as a sequence identifier. Protein with identical ordered strings of D, E, K, R were grouped together. A protein was considered unique by order if the reduced sequence appeared only once in the proteome.

#### 3. Effective reduction of noise using multiple probes

An important aspect of nanopore electrometry is the potential ability to perform simultaneous measurements of fields using multiple sensors at different positions. Fig.S1 shows representative raw signals of electric fields and currents calculated in this work. Importantly, we notice that averaging the signals over multiple probes leads to noise reduction.

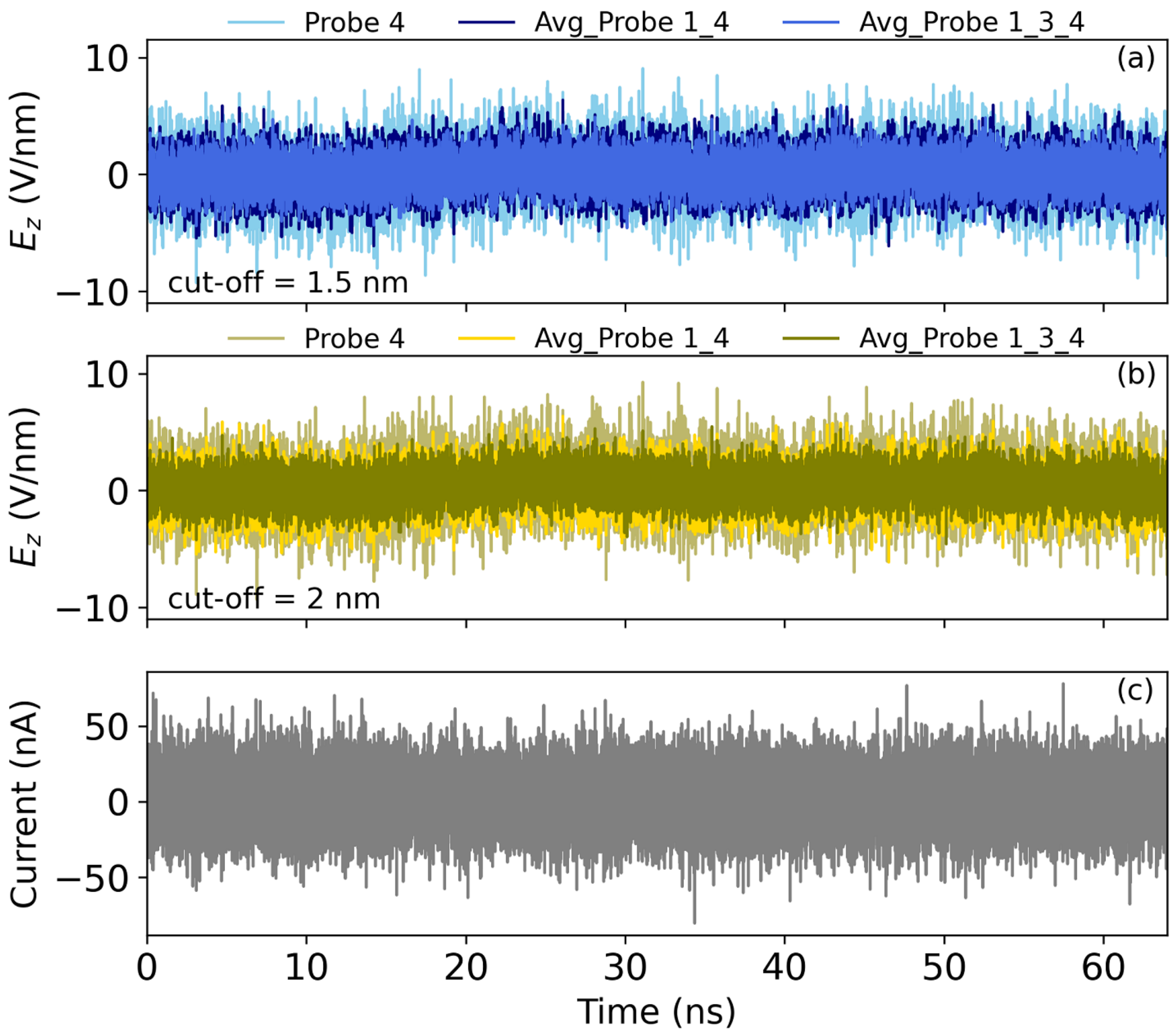


**Figure S1:** Raw signals of electric fields and currents

#### 4. Representative Processed Traces for all peptides

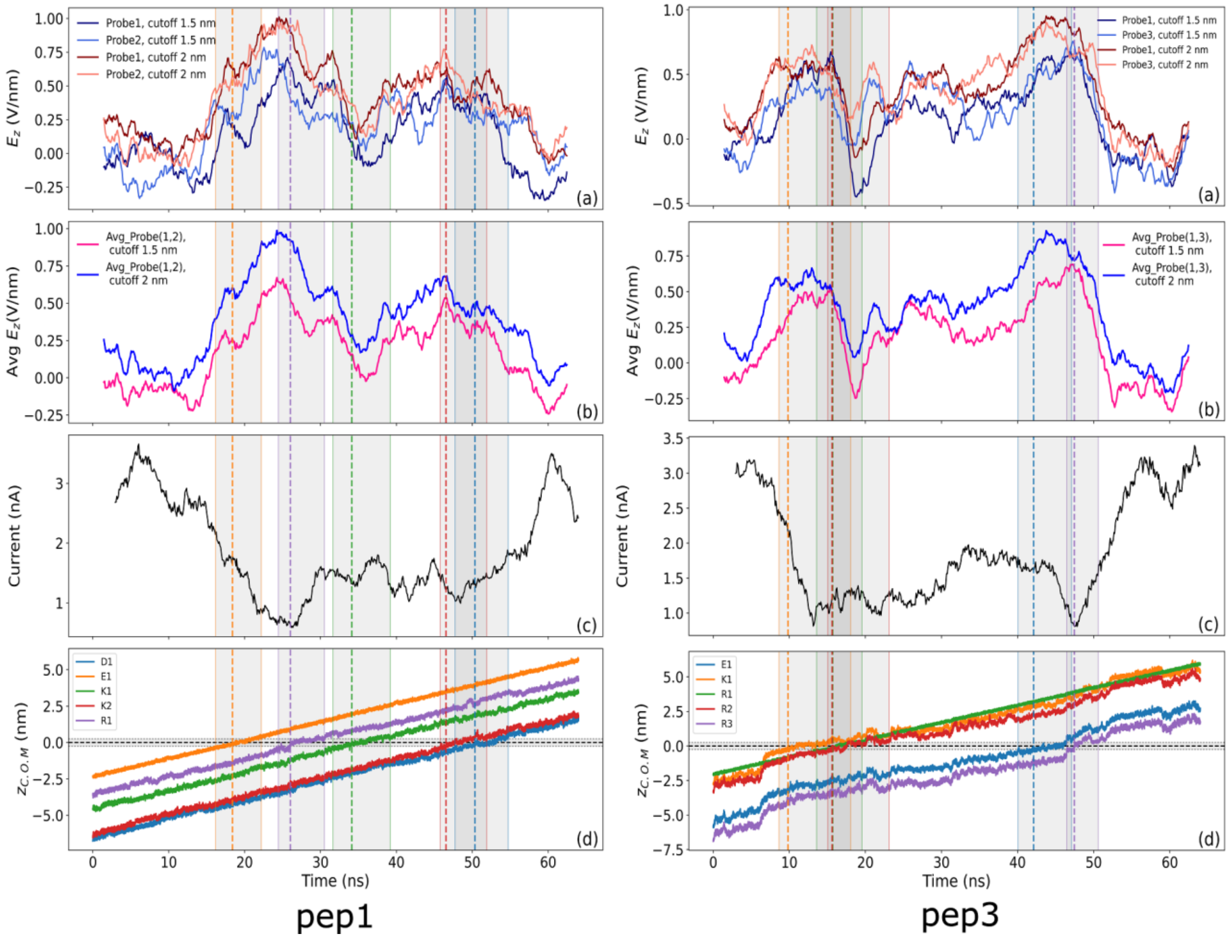


**Figure S2:** Time dependent signals while pep1, pep3 translocate through the pore. (a) Electric field traces captured by different probes. (b) Electric field trace after averaging over data obtained from two probes. (c) Ionic current signature. (d) Position of the charged residues with respect to the pore. The gray bands indicate their passage through the pore within $|Z^{res}_{c.o.m.} - Z_{pore}| = 0.25\ nm$. The band edges are color matched with the residues. The dotted vertical lines represent $|Z^{res}_{c.o.m.} - Z_{pore}| = 0$. The residue label, for example, K2 represents residue K that is arriving at the pore for the second time, as per the sequence of the peptide.

Both pep1 and pep3 has five charged amino acids in their sequence. However, the charged residues and their positional distributions are different (Fig.S2, panel d for both peptides). The corresponding traces are also significantly different. In panel a, b, we compare the electric field traces subjected to the sensing regions with cutoff 1.5nm and 2nm, respectively. No significant qualitative deterioration is observed. In the following, we provide the translocation signatures of pep2 and pep4. Each of them has two charged amino acids with different spatial distribution.

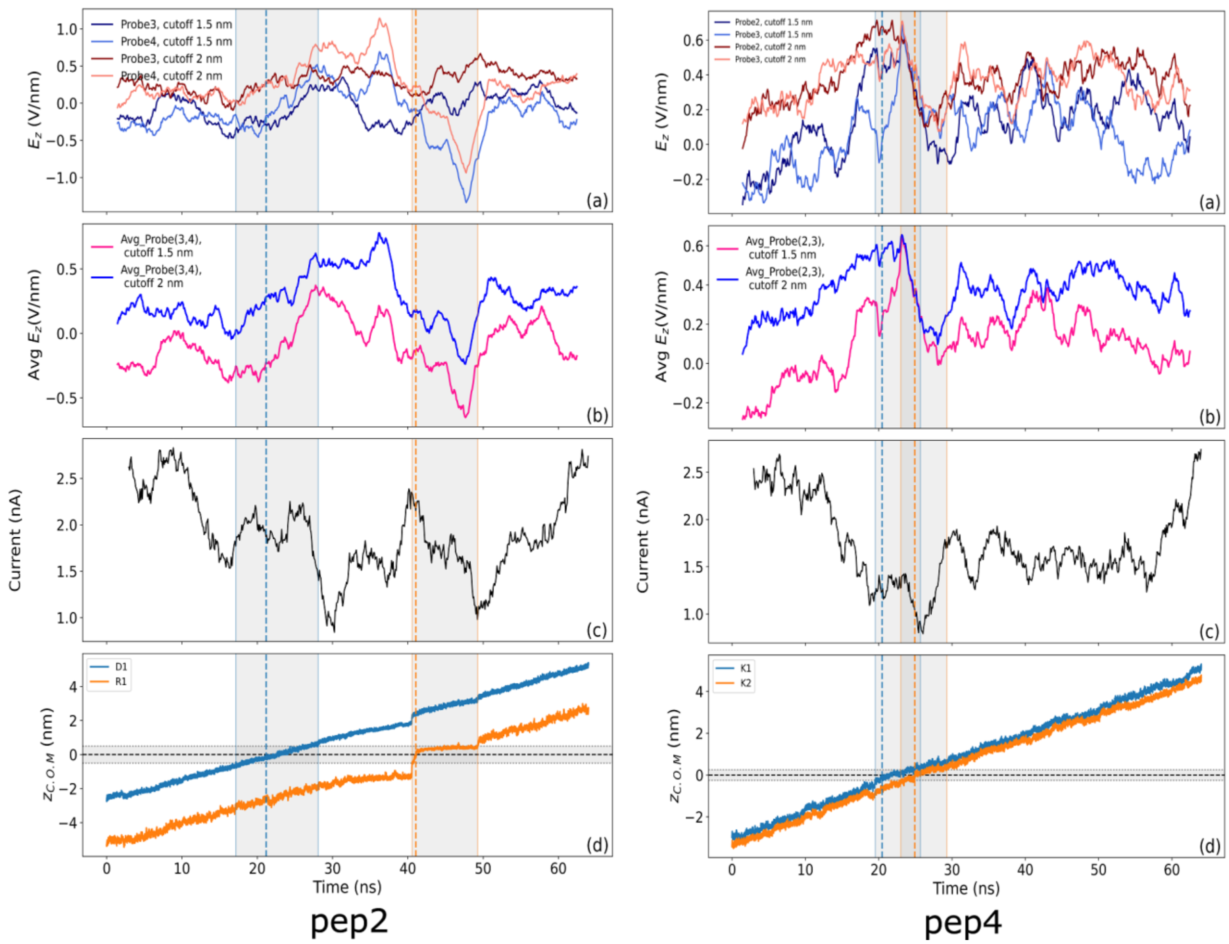


**Figure S3:** Time dependent signals while pep2, pep4 translocate through the pore. (a) Electric field traces captured by different probes. (b) Electric field trace after averaging over data obtained from two probes. (c) Ionic current signature. (d) Position of the charged residues with respect to the pore.

#### 5. Peak resolution comparison among the field and current histograms

The separation between adjacent modes of each Gaussian mixture model (GMM) in Figure 3, 4 was quantified using a peak resolution metric calculated from the fitted probability density function. The GMM component means were ordered in ascending value, and for each pair of consecutive components the minimum probability density between the two-component means was identified as the inter-peak valley. The peak resolution was then calculated as

$$R = 1 - \frac{2y_{valley}}{y_{peak1} + y_{peak2}}$$

where $y_{peak1}$ and $y_{peak2}$ are the probability densities at the two consecutive component means, and $y_{valley}$ is the minimum probability density between them. R=1 means the peaks are perfectly resolved, whereas R=0 means not resolved at all. Pairwise peak resolutions were calculated for every adjacent mode pair, while unimodal distributions were not assigned a peak resolution. The mean peak resolution, obtained by averaging all adjacent pair resolutions within a distribution, was used as a single descriptor of overall modal separability. For electric-field sensing, the highest mean peak resolution among the four probes was compared with the corresponding ionic-current value for each peptide.

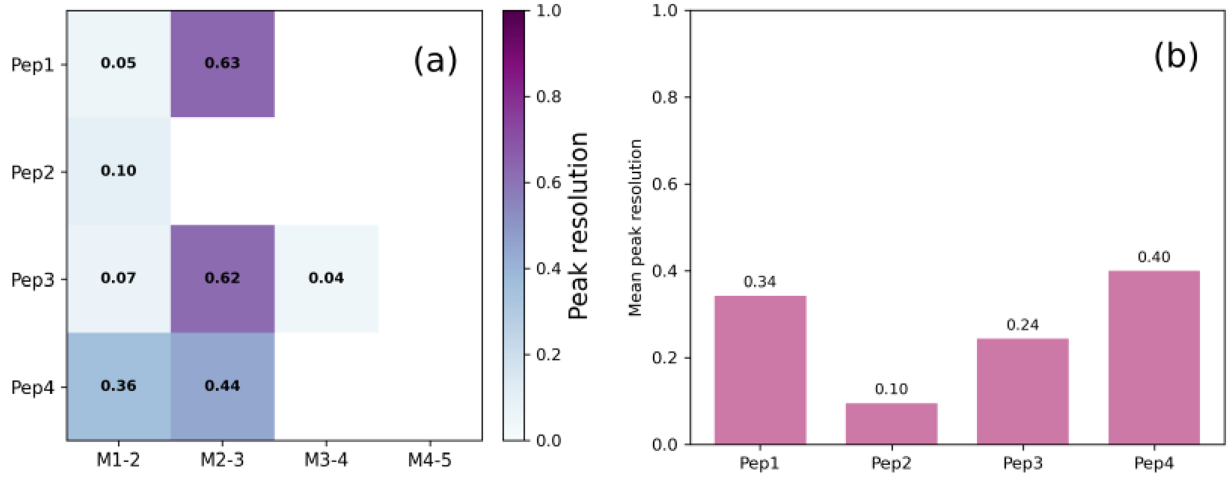


**Figure S4:** (a) Pairwise peak resolutions were calculated for every adjacent mode pair in current distribution (Fig.4). (b) The mean peak resolution, obtained by averaging all adjacent pair resolutions in a, was used to quantify overall modal separability.

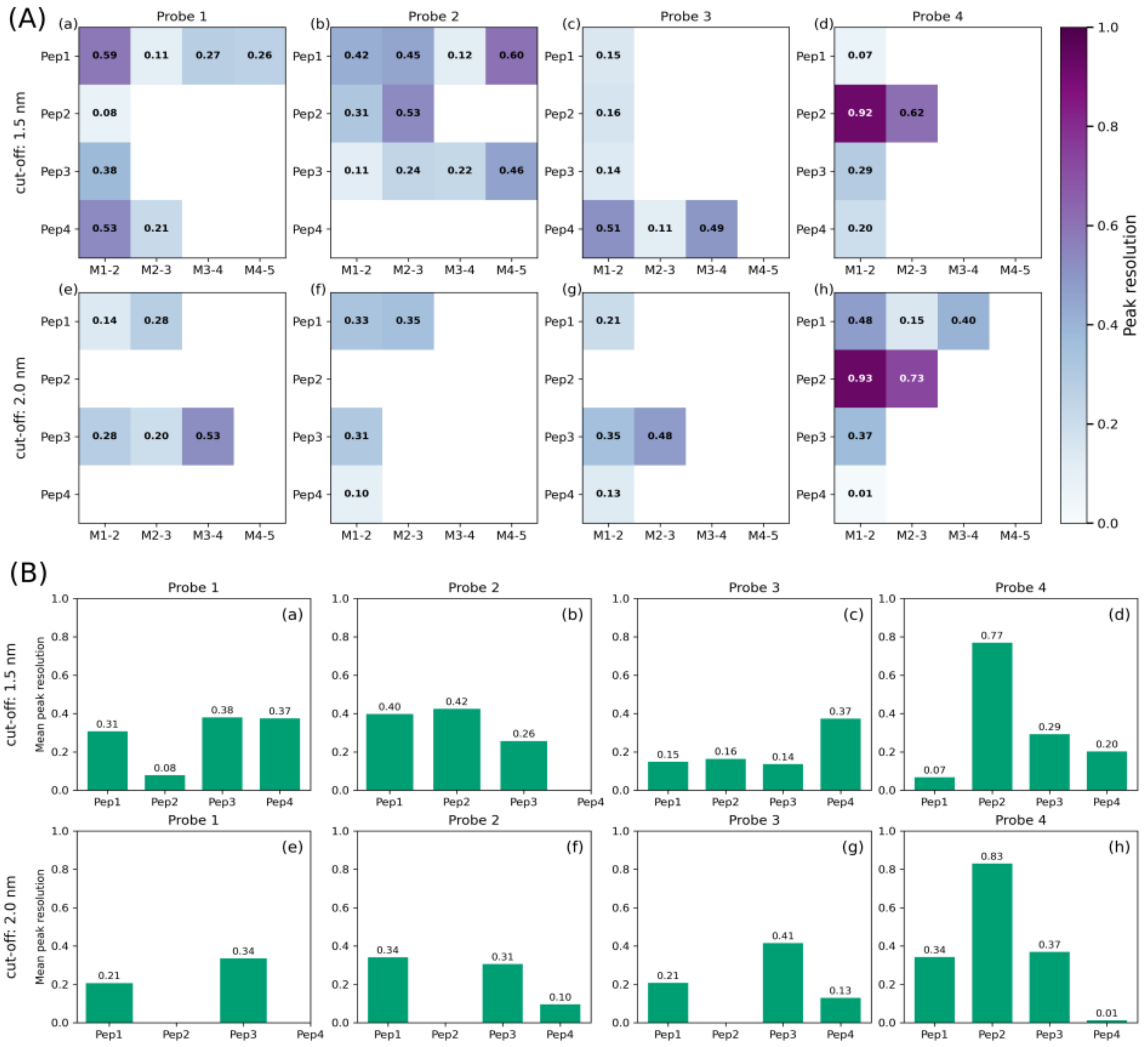


**Figure S5:** (A) Pairwise peak resolutions were calculated for every adjacent mode pair in current distribution (Fig.3). The gaps correspond to unimodal distributions as they were not assigned a peak resolution. (B) The mean peak resolution, obtained by averaging all adjacent pair resolutions in A, was used to quantify overall modal separability.

In Figure S6, the highest mean peak resolution for electric field among the four probes (Figure S5 B) is compared with the mean peak resolution of ionic-current value (Figure S4 b) for each peptide. The comparison indicates that mean peak resolution for field is higher than current.

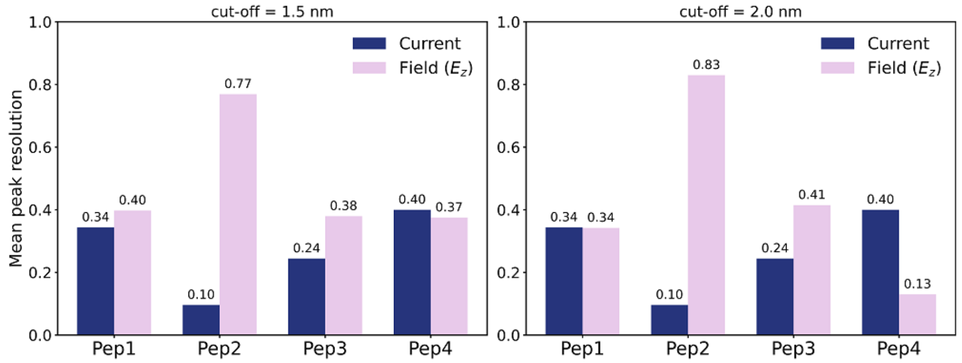


**Figure S6:** Mean peak resolution comparison between field and current

### 6. DTW distances across different probes

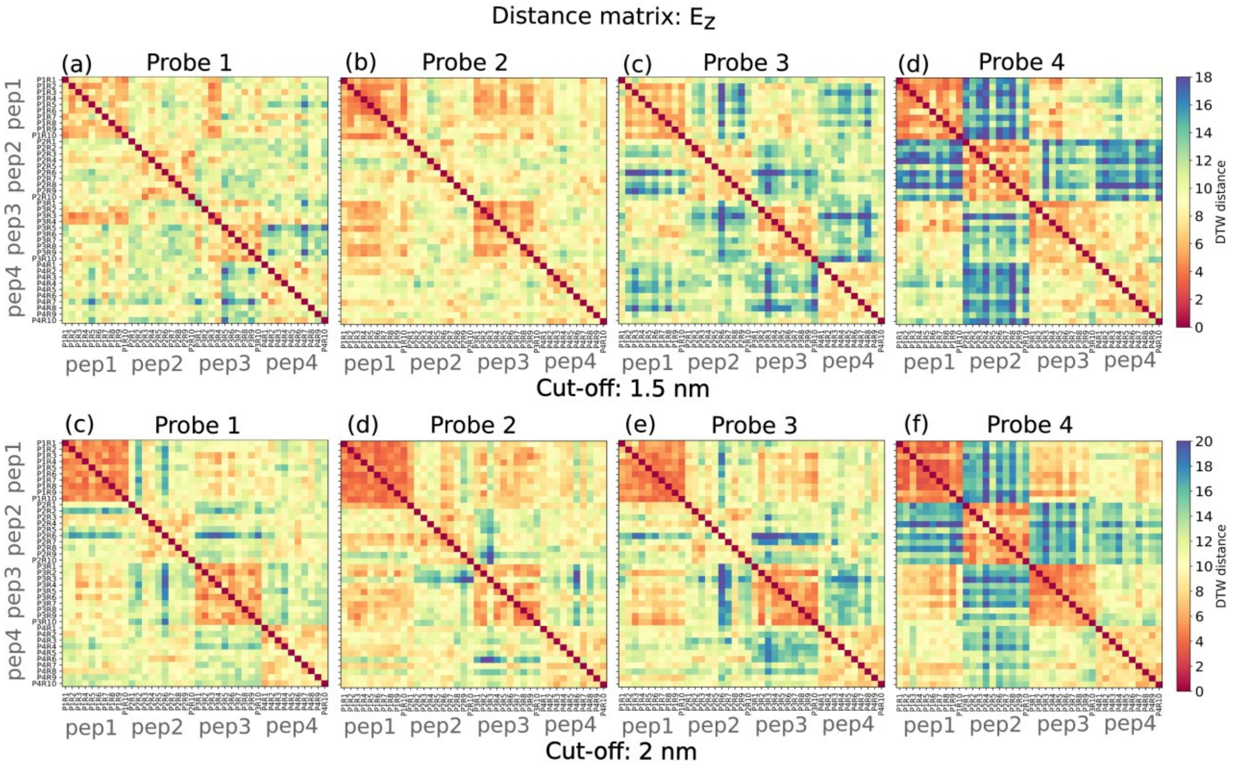


**Figure S7:** DTW distance matrices calculated among the signals from each probe. The top row (a, b, c, d) corresponds to the sensing region with 1.5nm cutoff, whereas the bottom row (c, d, e, f) corresponds to the sensing region with 2nm cutoff.

Figure S6 illustrates a fingerprint of peptide-specific electric field distinction among independent translocations realized by different probes.

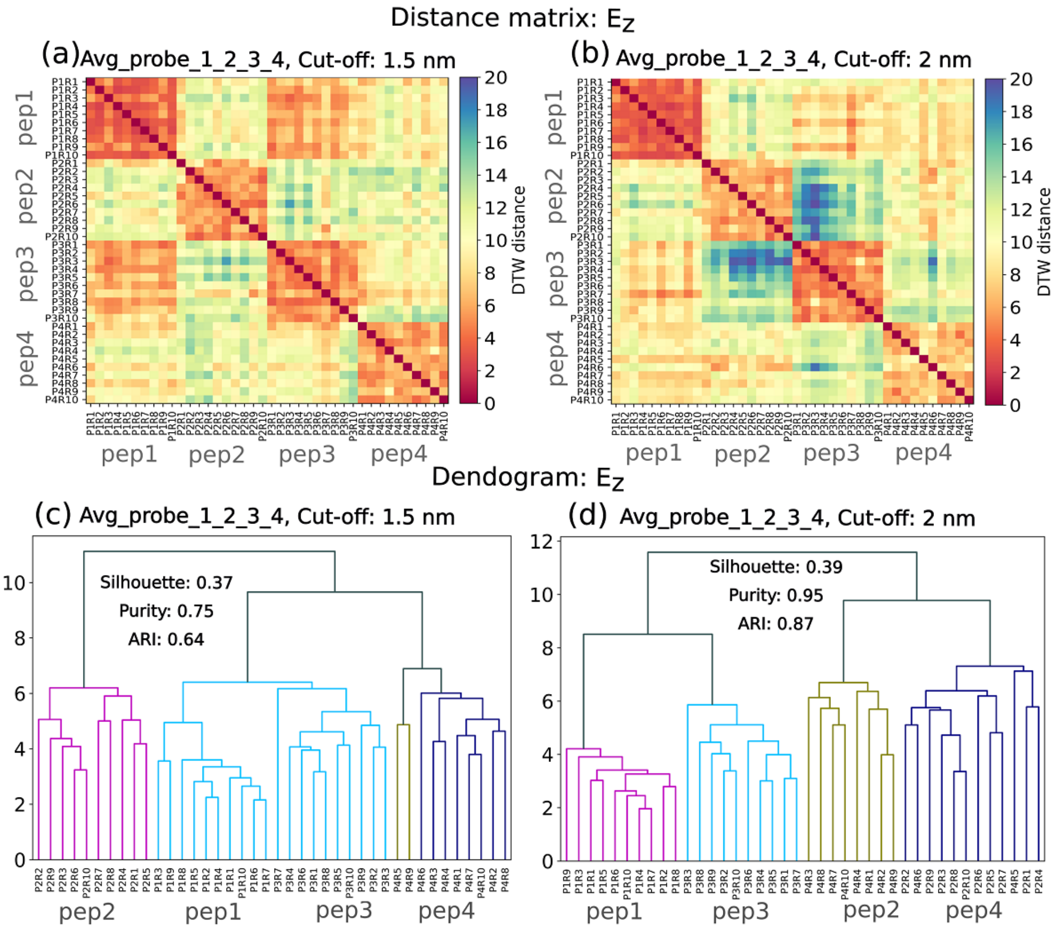


**Figure S8:** Dynamic time warping (DTW) distance matrices computed from ten independent translocation replicas of each peptide using the averaged z-component of the electric field from the average of all probes within sensing volume corresponding to radial cut-off of (a) 1.5nm and (b) 2nm. Hierarchical clustering of the same DTW distances for (c) electric field within 1.5nm sensing radius, (d) electric field within 2nm sensing radius.

Figure S8 demonstrates how the signal information is enriched when instead of using a single probe, all the probe signals are averaged. Compared to Figure S7, the distance matrices in Figure S8 clearly show improved peptide specific ordering, which was previously discussed in context of Figure 5.

To assess whether the clustering reported in the main text depends on any individual translocation trajectory, a leave-one-replicate-out validation was performed on each of the three dynamic time warping distance matrices, corresponding to the electric field at the 1.5 nm and 2 nm sensing volumes and to the ionic current. The ten replicas of every peptide are indexed identically; in each of ten folds the replica of the corresponding index was removed from all four peptides, leaving thirty-six trajectories, and the four-cluster solution was recomputed on the retained submatrix under the same average-linkage procedure used for the full data set. The adjusted Rand index, purity, and silhouette coefficient were evaluated against the known peptide labels for each fold, and the minimum and maximum of each metric across the ten folds are reported as its leave-one-replicate-out range. The procedure is deterministic and enumerates all ten folds exhaustively, requires no additional simulation, and was carried out with the same analysis script used for the confidence-interval estimation, so that the ranges reported below are exactly reproducible. Under this validation the electric-field clustering was found to be stable. At the 1.5 nm sensing volume the adjusted Rand index, purity, and silhouette coefficient remained within [0.63, 1.00], [0.75, 1.00], and [0.32, 0.43], respectively, and at the 2 nm sensing volume within [0.85, 1.00], [0.94, 1.00], and [0.37, 0.41]. For the ionic current the corresponding ranges were [0.12, 0.27], [0.47, 0.56], and [0.13, 0.20]. No single replicate was therefore responsible for the observed separation. Consistent with the confidence-interval analysis, the perfect adjusted Rand index obtained at the 1.5 nm sensing volume is the more sensitive to replicate removal, its lower bound of 0.63 lying below the value of 0.85 obtained at 2 nm; nonetheless, for all three metrics the electric-field ranges remain disjoint from those of the ionic current, so that the superior separability of the field-based readout is preserved throughout the validation.
